# DnaJB6 is a RanGTP-regulated protein involved in dynein-dependent microtubule organization during mitosis

**DOI:** 10.1101/536045

**Authors:** Rosas-Salvans M., Isabelle Vernos

## Abstract

Bipolar spindle organization is essential for the faithful segregation of chromosomes during cell division. This organization relies on the collective activities of motor proteins. The minus-end directed dynein motor complex generates spindle inward forces and plays a major role in spindle pole focusing. The dynactin complex regulates many dynein functions increasing its processivity and force production.

Here we show that DnaJB6 is a novel RanGTP regulated protein. It interacts with dynactin p150Glued in a RanGTP-dependent manner specifically in M-phase and promotes spindle pole focusing and dynein force generation. Our data suggest a novel mechanism by which RanGTP regulates dynein activity during M-phase.

**Summary statement:** We describe DnaJB6 as a novel RanGTP-regulated protein important for spindle assembly. Our data suggest that RanGTP regulates dynein-dependent inward spindle force generation and pole focusing through DnaJB6

## Introduction

Spindle assembly involves the organization of microtubules into two antiparallel arrays with interdigitating plus-ends and two focused minus-ends forming the spindle poles. This organization is highly dependent on the collective activities of microtubule-dependent motor proteins. The plus-end directed tetrameric motor Eg5 generates outward forces that together with other mechanisms separate the spindle poles (Blangy et al., 1995; Kapitein et al., 2005; Kashina et al., 1996; Kashina et al., 1997; Whitehead and Rattner, 1998). Indeed interfering with Eg5 activity results in the assembly of monopolar spindles with unseparated centrosomes (Mayer et al., 1999). Interestingly, impairing dynein activity rescues spindle bipolarity in Eg5 inhibited cells (Ferenz et al., 2009; Mitchison et al., 2005; Raaijmakers et al., 2013; Tanenbaum et al., 2008). This suggests that a fine balance between outward forces generated by Eg5 and inward forces generated by dynein defines spindle length and bipolarity (Ferenz et al., 2009; Florian and Mayer, 2012; Gaglio et al., 1997; Gaglio et al., 1996; Heald et al., 1996; Tanenbaum et al., 2008; Walczak et al., 1998).

In addition to force generation within the spindle, dynein is also essential for focusing microtubule minus-ends into the spindle poles (Echeverri et al., 1996; Gaglio et al., 1996; Heald et al., 1996; Mitchison et al., 2005; Tan et al., 2018; Walczak et al., 1998; Wittmann and Hyman, 1999). This process involves NuMA that recruits the dynein-dynactin complex to the microtubule minus-ends, promoting the transport and focusing of the microtubule minus-ends at the spindle pole (Compton and Cleveland, 1993; Gaglio et al., 1995; Haren et al., 2009; Hueschen et al., 2017; Khodjakov et al., 2003; Kisurina-Evgenieva et al., 2004; Merdes et al., 2000; Merdes et al., 1996; Silk et al., 2009).

During M-phase RanGTP triggers the nucleation, stabilization and organization of microtubules in the vicinity of the chromosomes playing an essential role in bipolar spindle assembly (Carazo-Salas et al., 1999; Cavazza and Vernos, 2015; Clarke and Zhang, 2008). The general mechanism involves the release of a specific group of NLS containing proteins from inhibitory interactions with importins (Gorlich et al., 1995; Gorlich et al., 1996). Several RanGTP regulated proteins have been identified and shown to play essential roles in spindle assembly (Cavazza and Vernos, 2015). Recently we used a mass spectrometry based proteomic approach to identify the full proteome associated with RanGTP induced microtubules in Xenopus egg extracts and validated a few selected proteins with no reported function in mitosis (Rosas-Salvans et al., 2018). One of them DnaJB6 (also known as MRJ, HSJ2 or MSJ1) had also been identified in mitotic spindle proteomes (Bonner et al., 2011; Rao et al., 2016; Sauer et al., 2005) although its function in mitosis had not been addressed. DnaJB6 is an HSP40 family protein with two alternatively spliced isoforms (in human). These two isoforms share the first 239 amino acids but differ at their C-terminus. The short isoform (DnaJB6-S, 241aa) is cytoplasmic whereas the long isoform (DnaJB6-L, 326aa) that has a nuclear localization signal at its C-terminus (Mitra et al., 2008) was shown to localize to the nucleus (Cheng et al., 2008; Mitra et al., 2008). The co-chaperon activity of the short isoform has been demonstrated in vitro (Chuang et al., 2002). Additionally, DnaJB6 prevents huntingtin aggregation *in vitro* independently of the DnaJ domain (and HSP70), although *in vivo* this domain seems to be required (Chuang et al., 2002; Hageman et al., 2010). DnaJB6-L expression is associated to suppression of tumourogenesis and metastasis in breast cancer (Meng et al., 2016). Interestingly, DnaJB6 is highly expressed in human testis, ovary, liver and placenta (Seki et al., 1999) and its expression is increased in mitosis in HeLa cells (Dey et al., 2009; Seki et al., 1999). We previously showed that DnaJB6 localizes to the spindle poles and its silencing affects microtubule aster formation in cells undergoing microtubule regrowth, suggesting that it is a novel RanGTP regulated SAF (Spindle Assembly Factor) (Rosas-Salvans et al., 2018).

Here we show that DnaJB6 co-pulls down the dynactin complex component p150Glued in a RanGTP-dependent manner. Moreover, DnaJB6 is required for inward force generation during bipolar spindle assembly and spindle pole focusing, both dependent on dynein activity. Our data suggest that dynein may be regulated during mitosis through a novel mechanism involving DnaJB6 and RanGTP.

## Results

### DnaJB6 is a RanGTP regulated protein involved in spindle assembly

We first directly tested whether DnaJB6-L (that contains an NLS) could be regulated by RanGTP during mitosis using the Xenopus egg extract system. GST-xDnaJB6-L coated beads were incubated in CSF-arrested egg extract supplemented or not with RanGTP. Western blot (WB) analysis of the proteins associated with the beads showed that importin-β was specifically pulled down by GST-xDnaJB6-L in the absence but not in the presence of RanGTP (Fig 1A). This result suggests that DnaJB6 may be regulated by RanGTP during mitosis like other SAFs. The high degree of conservation of the NLS sequence of the Xenopus and human DnaJB6 proteins (Fig 1B) suggests that the interaction with importin-β may be conserved in human.

**FIGURE 1.**
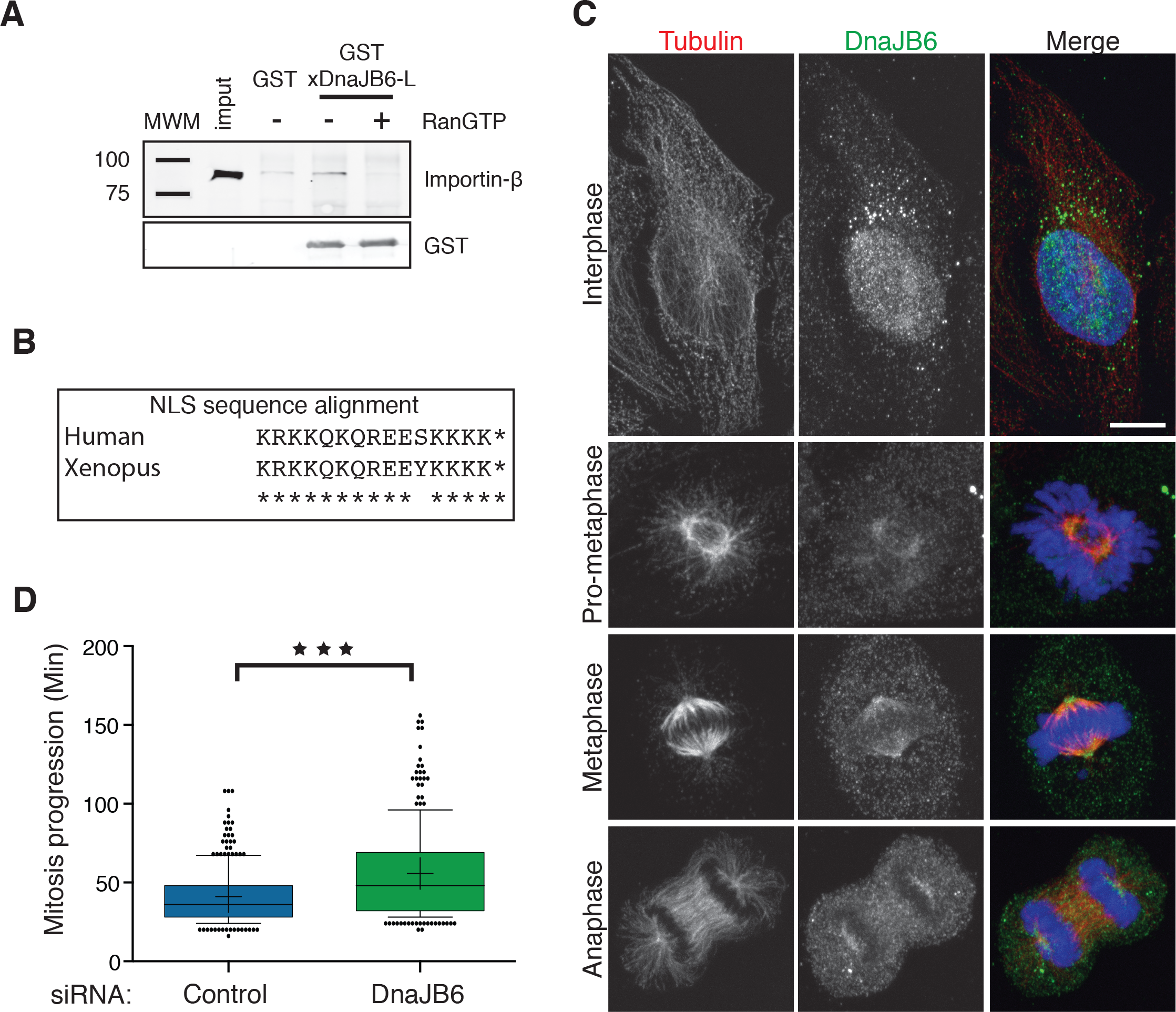
DnaJB6 is a RanGTP regulated protein required for mitotic progression. A. Western blot analysis of a GST-xDnaJB6 pulldown experiment in Xenopus egg extract. Importin-β associates with GST-xDnaJB6 and is realeased by addition of RanGTP to the extract (top). The lower panel shows that similar levels of GST-xDnaJB6 were used for pull downs in the presence or absence of RanGTP.
B. Amino acid sequence alignment of the putative NLS in human (top) and Xenopus (bottom) DanJB6 long isoforms.
C. Immunofluorescence images of HeLa cells showing the localization of DnaJB6 in different cell cycle stages. DnaJB6 accumulates in the nucleus during interphase and localizes to the spindle during mitosis with an accumulation at the spindle poles in metaphase. Tubulin is shown in red, DnaJB6 in green and DNA in blue. Scale bars, 10 μm.
D. Box and whiskers plot showing the time from the NEB to anaphase onset for control and DnaJB6 silenced cells. Boxes show the upper and lower quartiles (25-75%) with a line at the median, whiskers extend from the10th to the 90th percentile and dots correspond to outliers. The + indicates the mean values (41.1 for control and 55.7 minutes for DnaJB6 silenced cells). Data from three independent experiments in which a total of 281 control and 250 DanJB6 silenced cells were analyzed. Three asterisks indicate P <0.0001 (Mann-Whitney test).

We then used immunofluorescence microscopy to characterize the subcellular localization of DnaJB6 during the cell cycle. As previously described (Cheng et al., 2008; Mitra et al., 2008), DnaJB6 was predominantly in the nucleus during interphase as well as in the cytoplasm. Interestingly, in mitotic cells DnaJB6 was associated with the spindle microtubules with an accumulation at the spindle poles (Fig 1C).

To determine whether DnaJB6 has any role during mitosis we then used Time-lapse fluorescence microscopy to monitor mitosis progression in control and DnaJB6 silenced cells using a stable HeLa cell line expressing H2B–eGFP/ α-tubulin–mRFP. DnaJB6 silencing induced a delay in mitosis, from nuclear envelope breakdown to anaphase onset (Fig 1D). Indeed, these cells needed 35.5% more time than control cells to reach anaphase onset. To determine the causes for this delay we examined fixed control and DnaJB6 silenced HeLa cells by immunofluorescence microscopy (silencing efficiency shown in Fig S1A). The quantification of the distribution of the mitotic phases showed a significant decrease of DnaJB6 silenced cells in metaphase. Concomitantly the percentage of cells with spindle defects increased significantly and the percentage of pro-metaphase also increased although it was not statistically significant (Fig 2A). Altogether, these results suggested that DnaJB6 is required for spindle assembly and mitosis progression.

**FIGURE 2.**
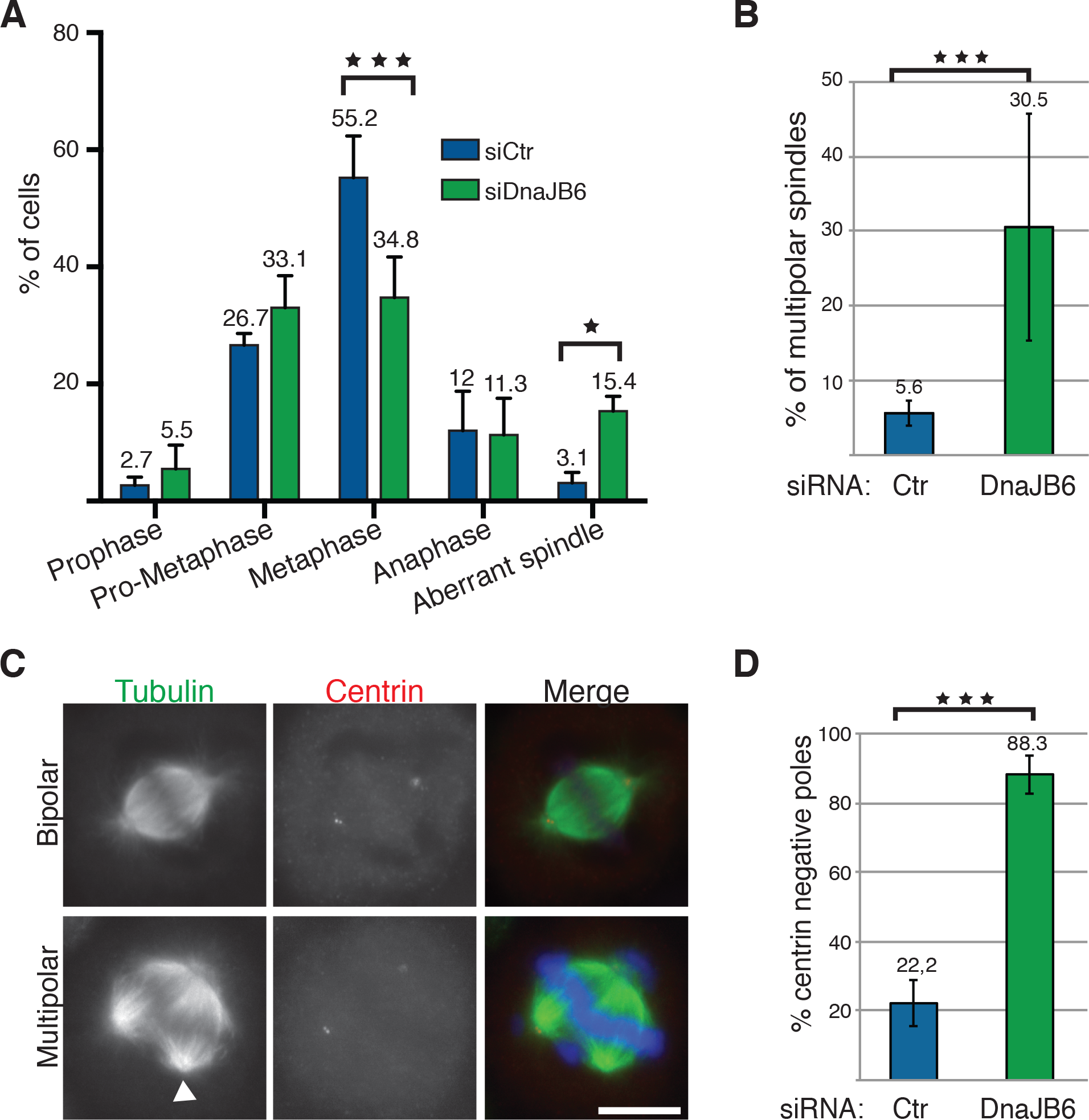
DnaJB6 is required for bipolar spindle assembly. A. Bars graph showing the distribution of mitotic phases in fixed control (blue) or DnaJB6 silenced HeLa cells (green). The percentage for each category is shown at the top of the columns. Monopolar and multipolar spindles as well as more disorganized structures were included into the ‘Aberrant spindle’ category. Data from three independent experiments in which a total of 454 control and 498 DnaJB6 silenced cells were analyzed. One asterisk indicate P<0.05, Three asterisks indicate P<0.0001 (ANOVA test).
B. Quantification of the multipolar spindles in control (blue) and DnaJB6 (green) silenced cells. Data from three independent experiments in which a total of 765 control and 778 DnaJB6 silenced cells were analyzed. Three asterisks indicate P <0.0001 (Fisher’s exact test).
C. Immunofluorescence images of DnaJB6 silenced HeLa cells showing tubulin (green), centrin (red) and DNA (blue). A bipolar and a multipolar spindle are shown. The arrow points to a centrin negative pole. Scale bar, 10μm.
D. Quantification of multipolar spindles with poles negative for centrin in control (blue) and DnaJB6 (green) silenced HeLa cells. The percentages are indicated on top of the columns. Data from two independent experiments in which a total of 126 control and 116 DnaJB6 silenced cells were analyzed. Three asterisks indicate P<0.0001 (ANOVA test).

### DnaJB6 is essential for microtubule organization during mitosis

To address the role of DnaJB6 in spindle assembly, we first examined more carefully the spindle morphology defects associated with DnaJB6 silencing. The major phenotypes were associated with microtubule organization defects such as multipolar spindles (Fig 2B) or the presence of ectopic microtubule clusters (Fig S1B). Indeed DnaJB6 silenced cells showed a significant increase of multipolar spindles compared to control cells (30.5% versus 5.6%) (Fig 2B). This phenotype was not associated to the amplification of the number of centrosomes in the silenced cells as monitored with anti-centrin antibodies. Indeed, 88% of the multipolar spindle assembled in DnaJB6 silenced cells had only two centrin positive poles whereas 78% of the few multipolar spindles assembled in control cells had centrin signal at all the poles (Fig 2C,D). These results suggested that DnaJB6 is required for microtubule organization during mitosis. To gain further support for this idea we monitored microtubule organization in control and DnaJB6 silenced cells undergoing regrowth after nocodazole washout. Cells were fixed at different time points after nocodazole washout and processed for immunofluorescence (Fig 3A). The efficiency of bipolar spindle organization was measured by quantifying the number of microtubule asters at the different time points (Fig S1C). In most control cells, the number of microtubule asters was initially high due to the activity of the centrosomal and chromosomal pathways(Meunier and Vernos, 2011) and reduced over time to finally only two microtubule asters defining the poles of the bipolar spindle. In contrast, the percentage of cells with only two microtubule asters was significantly reduced in DnaJB6 silenced cells at all the time points examined (Fig 3B) and the difference with the control increased with time. At 60 minutes after nocodazole washout 57.7% of the control cells had two asters organized into bipolar spindle, whereas this percentage was reduced to 24.8% in DnaJB6 silenced cells (Fig 3C).

**FIGURE 3.**
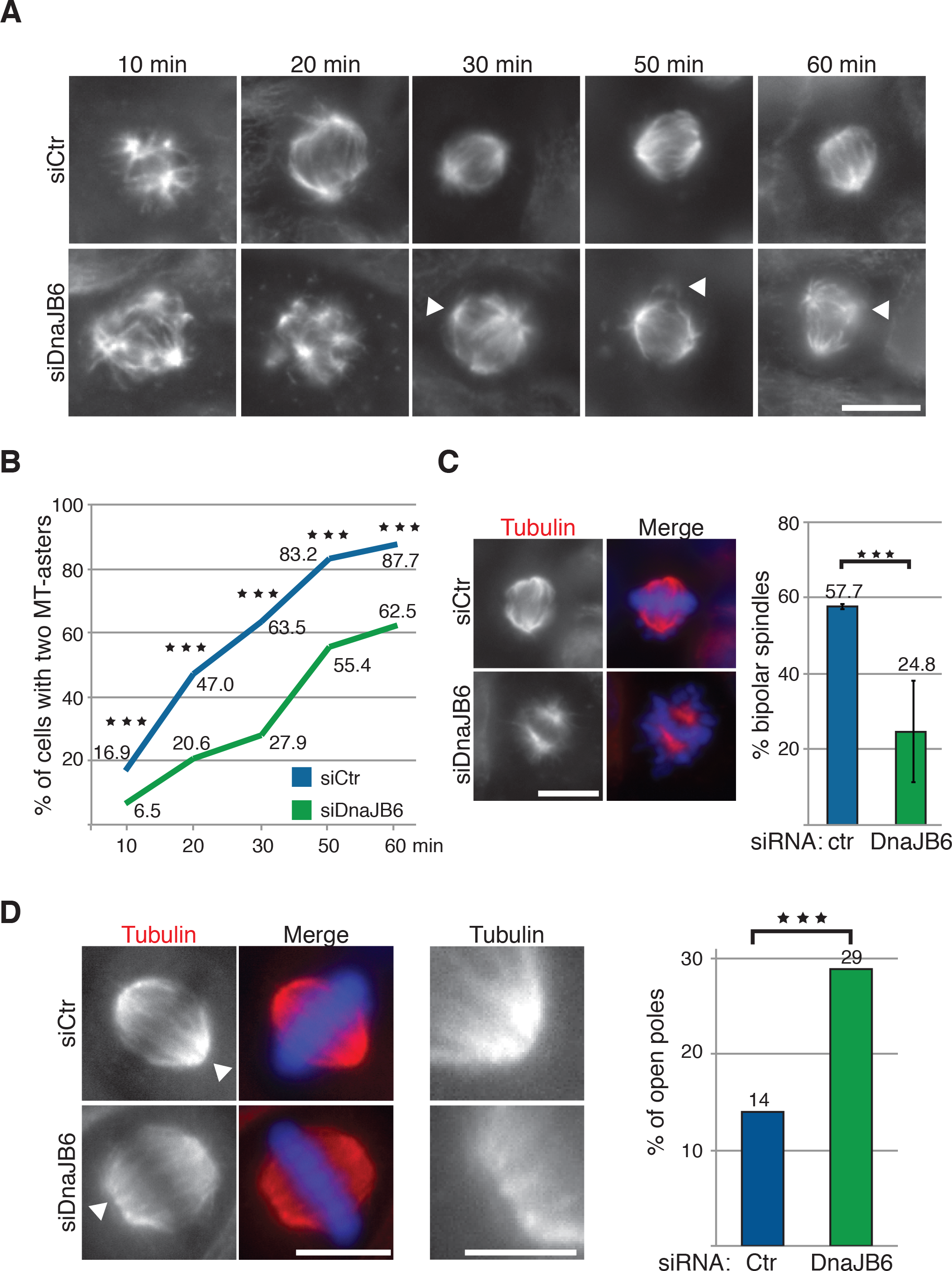
DnaJB6 is involved in spindle microtubules organization. A. Immunofluorescence images of mitotic control and DnaJB6 silenced HeLa cells undergoing microtubule regrowth after nocodazole washout. Cells were fixed at the times indicated after nocodazole washout and processed for immunofluorescence to visualize the microtubules and DNA (not shown). The white arrows point to microtubule asters that fail to incorporate into the main spindle poles. Scale bar, 10μm.
B. Quantification of the percentage of cells with two asters at different time points after nocodazole washout as shown in A. The blue line correspond to the control cells and the green line to DnaJB6 silenced cells. Data from three independent experiments in which more than 300 cells were recorded for each time point. Three asterisks indicate P<0.001 (Fisher’s exact test).
C. Representative immunofluorescence image of a HeLa cell fixed at 60min after nocodazole washout with two microtubule asters that do or do not organize a bipolar spindle. In the merge, tubulin is in red and DNA in blue. Scale bar, 10μm. Quantification of the percentage of cells having two microtubule asters organized into a bipolar spindle 60 minutes after nocodazole washout. Data from two independent experiments in which 503 control (blue) and 638 DnaJB6 silenced cells (green) were analyzed. The exact percentages are indicated on top of each bar. Three asterisks indicate P<0.001(Fisher’s exact test)
D. Immunofluorescence images of bipolar spindles in control and DnaJB6 silenced HeLa cells (Scale bar 10μm). Amplified view of the spindle poles (indicated with white arrows) are shown on the right (Scale bar 5μm). Microtubules are unfocused at the spindle poles in DnaJB6 silenced cell. Tubulin is shown in red and DNA in blue. Graph showing the percentage of unfocused poles in 100 bipolar spindles of control (blue) and DnaJB6 silenced cells (green). The three asterisks indicate P<0.001 (Fisher’s exact test).

The strong defects in microtubule aster clustering and the presence of centrin-negative extra spindle poles suggested that DnaJB6 could also be required for spindle pole focusing. We therefore decided to look more carefully at the morphology of the poles of bipolar spindles in the DnaJB6 silenced cells (Fig 3D). Indeed we found that 29% of the poles of the bipolar spindles were not tightly focused in these cells whereas this was only observed in 14% of the bipolar spindles assembled in control cells (Fig 3D).

Altogether, these results strongly suggest that DnaJB6 plays a role in microtubule organization in mitosis, more specifically in microtubule aster clustering and spindle pole focusing.

### DnaJB6 has a direct role in microtubule organization in M-phase

Since DnaJB6 is a co-chaperon, its absence could potentially lead to an accumulation of protein folding defects that could be related to the phenotypes observed during mitosis. To determine whether the phenotypes we observed in DnaJB6 silenced cells could be due to indirect effects on protein stability we used the Xenopus egg extract system. Cycled spindle assembly experiments were performed in control and DnaJB6 depleted egg extracts. In addition in rescue experiments, recombinant MBP or MBP-xDnaJB6-L were added to the depleted extracts as the extract was sent back into mitosis (Fig 4A and Fig S2). Samples were collected 60min after cycling the extract into mitosis and processed for immunofluorescence. Although bipolar spindles assembled with similar efficiency in control and DnaJB6 depleted extracts (data not shown), the absence of DnaJB6 resulted in spindle pole defects (Fig 4B). While 86.8% of the control spindles had well focused poles, spindles assembled in DnaJB6 depleted extracts had a statistically significant reduction of focused spindle poles to 64%. Indeed in depleted extracts there was a high proportion of open spindle poles sometimes even generating two sub-poles. This phenotype was rescued by addition of MBP-xDnaJB6-L to the depleted extract as 81% of the spindle poles were focused (Fig 4C). This showed that the spindle assembly and spindle pole focusing defects can be entirely attributed to the absence of DnaJB6 specifically during mitosis. Consistently, the addition of an excess of MBP-xDnaJB6-L to the egg extracts as they were cycled into mitosis resulted in an increase of the percentage of very tightly focused spindle poles (Fig 4D).

**FIGURE 4.**
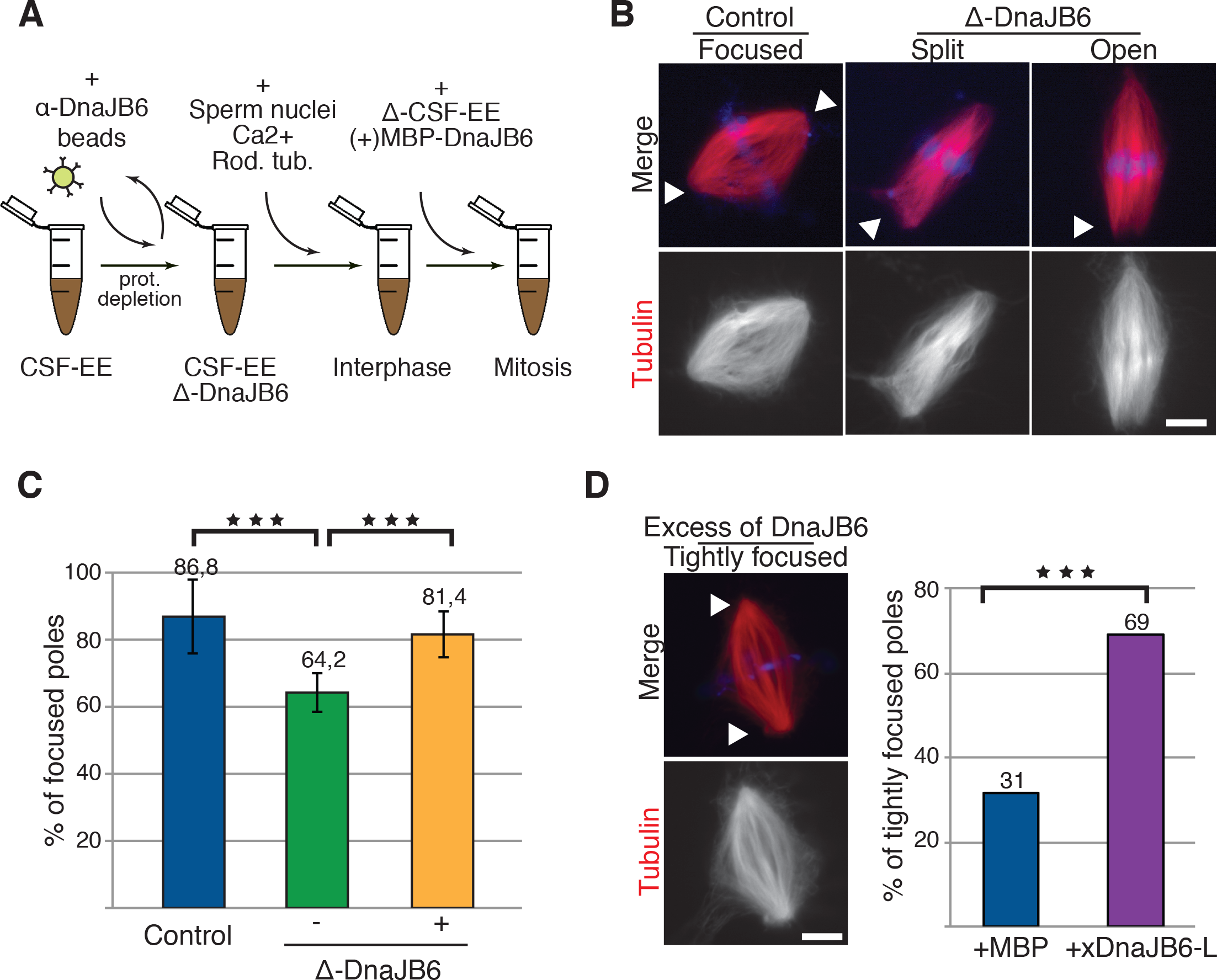
DnaJB6 depletion from Xenopus egg extracts induce spindle pole organization defects that are rescued by addition of recombinant protein. A. Schematic representation of the experimental procedures for spindle assembly experiments in Xenopus egg extracts.
B. Fluorescence images of spindles assembled in control or DnaJB6 depleted egg extracts. Most spindles assembled in control depleted extracts have focused spindle poles whereas many spindles assembled in DnaJB6 depleted extracts show spindle pole focusing defects defined as ‘Split poles’ or ‘Open poles’ as shown. The white arrows point at to the different types of spindle poles. In the merge, tubulin is shown in red and DNA in blue. Scale bar, 10μm.
C. Graph showing the quantification of focused spindle poles in control extracts (blue), DnaJB6 depleted extracts (green) and DnaJB6 depleted extracts containing MBP-xDnaJB6 (orange). Data from four independent experiments in which 703, 607 and 358 spindle poles were examined respectively. The three asterisks indicate P <0.001 (ANOVA test).
D. Fluorescence image of a spindle assembled in egg extract supplemented with an excess of DnaJB6 (1μM) showing tightly focused spindle poles (white arrows). Scale bar 10μm. Bars graph (at the right) shows the percentage of tightly focused poles in spindles assembled in control extracts supplemented with MBP (blue) and in extracts supplemented with MBP-xDnaJB6 (magenta). Data from three independent experiments in which 114 and 258 spindle poles were analyzed. Three asterisks indicate P<0.0001(Fisher exact test).

Together, these results indicate that DnaJB6 plays a direct role in microtubule focusing at the spindle poles during M-phase.

### DnaJB6 interacts with p150Glued in a RanGTP-dependent manner during mitosis and is required for its spindle localization

The defects in microtubule organization and spindle pole focusing associated to the absence of DnaJB6 were reminiscent of those induced by interfering with minus-end directed motor activities (Kardon and Vale, 2009; Raaijmakers and Medema, 2014). To test whether DnaJB6 may interact with minus ends motors we performed pull-down experiments in egg extracts using MBP-xDnaJB6-L or MBP coated beads in the presence or absence of RanGTP. Western blot analysis showed that the dynactin subunit p150Glued was specifically pulled down by MBP-xDnaJB6 but only in the presence of RanGTP (Fig 5A). These results suggested that DnaJB6 interacts with p150Glued in a RanGTP dependent manner during M-phase.

**FIGURE 5.**
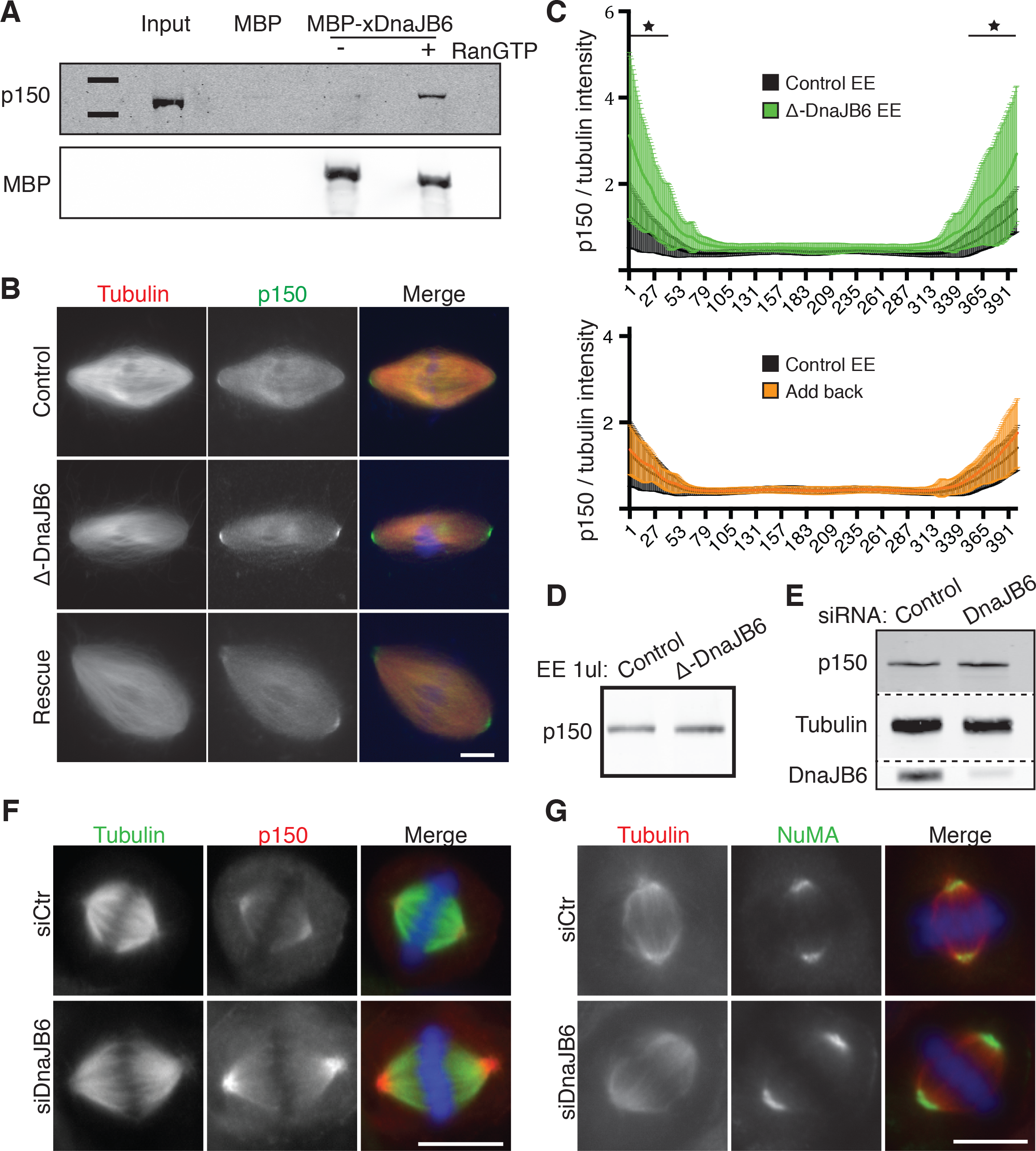
DnaJB6 interacts with p150 in a RanGTP dependent manner and is required for its spindle localization. A. Western blot analysis of a GST-xDnaJB6 pulldown experiment in Xenopus egg extract. MBP-xDnaJB6 or MBP coated Dynabeads were incubated in egg extracts in the presence or absence of RanGTP. P150Glued was specifically pulled down with MBP-xDnaJB6 from extracts containing RanGTP. The lower panel shows that similar amounts of MBP-xDnaJB6 was used for pull down in the presence or absence of RanGTP.
B. Immunofluorescence images showing P150Glued localization on bipolar spindles assembled in control extracts, DnaJB6 depleted extracts or DnaJB6 depleted extracts supplemented with recombinant MBP-xDnaJB6. p150Glued is shown in green, tubulin in red and DNA in blue. Scale bar, 10μm.
C. Graphical representation of the p150Glued relative fluorescence signal intensity along the spindle. Fluorescence intensities from several images were analyzed using FIJI by drawing a rectangle (of a conserved size) from pole to pole on each spindle. The p150Glued intensity values were normalized using the tubulin fluorescence intensity signal. The average values are shown as a line within the standard deviation. Values from spindle assembled in control extracts are in black (n=20), those from spindles assembled in DnaJB6 depleted extracts are in green (n=16) and those assembled in DnaJB6 depleted extracts containing MBP-xDnaJB6 (add back) are in orange (n=21). Metaphase spindles were selected randomly, excluding the spindles with pole focusing defects. A statistically significant accumulation of p150Glued at the spindle poles occurs in DnaJB6 depleted extracts (P<0.05, two tailed ANOVA test) and is recued by addition of recombinant MBP-xDnaJB6 to the depleted extract (lower graph).
D. Western blot analysis of control and DnaJB6 depleted egg extracts (1μl each) showing that the levels of p150Glued are similar.
E. Western blot analysis of control and DnaJB6 cell lysates (30μg of total protein) showing that the levels of p150Glued are similar (top). Total α-Tubulin is shown in the middle as protein loading control and DnaJB6 silencing efficiency is shown at the bottom.
F. Representative immunofluorescence images from control and DnaJB6 silenced HeLa cells showing the localization of p150Glued in metaphase spindles. A stronger p150Glued signal is observed at the spindle poles of DnaJB6 silenced cells. In the merge, tubulin is in green, p150Glued in red and DNA in blue. Scale bar 10μm.
G. Representative immunofluorescence images from control and DnaJB6 silenced HeLa cells showing the localization of NuMA in metaphase spindles. The intensity of NuMa signal is higher at the spindle poles in DnaJB6 silenced cells. In the merge, tubulin is in green, NuMA in red and DNA in blue. Scale bar 10μm.

To gain further insights into the interaction between DnaJB6 and p150Glued we then checked whether p150Glued spindle localization was affected in the absence of DnaJB6. Spindles assembled in control or DnaJB6 depleted extracts supplemented or not with MBP or MBP-xDnaJB6-L were processed for immunofluorescence analysis to monitor p150Glued localization. In control spindles p150Glued localized all along the spindle microtubules with some enrichment at the spindle poles. In the absence of DnaJB6, p150Glued accumulated at the spindle poles (Fig 5B, C). Western blot analysis showed that the general levels of p150Glued were similar in control and DnaJB6 depleted extracts (Fig 5D). Rescue experiments showed that p150Glued accumulation was reverted to control levels at the spindle poles upon addition of MBP-xDnaJB6-L to the depleted extract (Fig 5B, C). These results indicated that DnaJB6 plays a role in p150Glued spindle pole localization in egg extracts.

Interestingly, p150Glued also accumulated to the spindle poles in DnaJB6 silenced cells (Fig 5F and S3A). Consistently with the results in egg extract, no differences on the general levels of p150glued were detected by western blot analysis of control and DnaJB6 silenced cell lysates (Fig 5E). Since it has been recently shown that p150Glued is recruited to the microtubule minus-ends by NuMA in mitosis (Hueschen et al., 2017), we then checked if NuMA localization was altered in DnaJB6-silenced cells. Immunofluorescence analysis showed that NuMA was also significantly enriched at the spindle poles in DnaJB6 silenced cells (Fig 5G and S3B).

Altogether, our data indicates that DnaJB6 interacts with p150Glued in a RanGTP-dependent manner during mitosis. Moreover, this interaction may be important for p150Glued and its targeting partner NuMa localization to the spindle poles.

### DnaJB6 increases the stability of dynactin specifically in mitosis

Since DnaJB6 is a co-chaperon, we reasoned that its interaction with p150Glued could have a role in the assembly or stabilization of the dynactin complex. To test this idea we performed sucrose density gradient analysis of cell lysates in the presence of the chaotropic agent potassium iodide (KI) previously shown to promote the dissociation of the dynein complex (King et al., 2002). Control and DnaJB6 silenced HeLa cells were synchronized in interphase and mitosis and the cell lysates incubated with KI before loading on top of a 8-20% sucrose density gradient. In control cells, western blot analysis showed that p150Glued migrated to lighter fractions of the gradient in the presence of KI. This indicated that KI induced the dissociation of the p150Glued complex both in interphase and mitosis, as described for the dynein complex. DnaJB6 silencing, did not change the dissociation pattern of the p150Glued complex in interphase. In contrast, it promoted a shift of p150Glued to lower sucrose concentrations in mitosis (Fig 6A). These results suggest that DnaJB6 is required for the stability of a p150Glued complex specifically in mitosis. This complex may be the dynactin complex itself or a larger complex including dynein.

**Figure 6.**
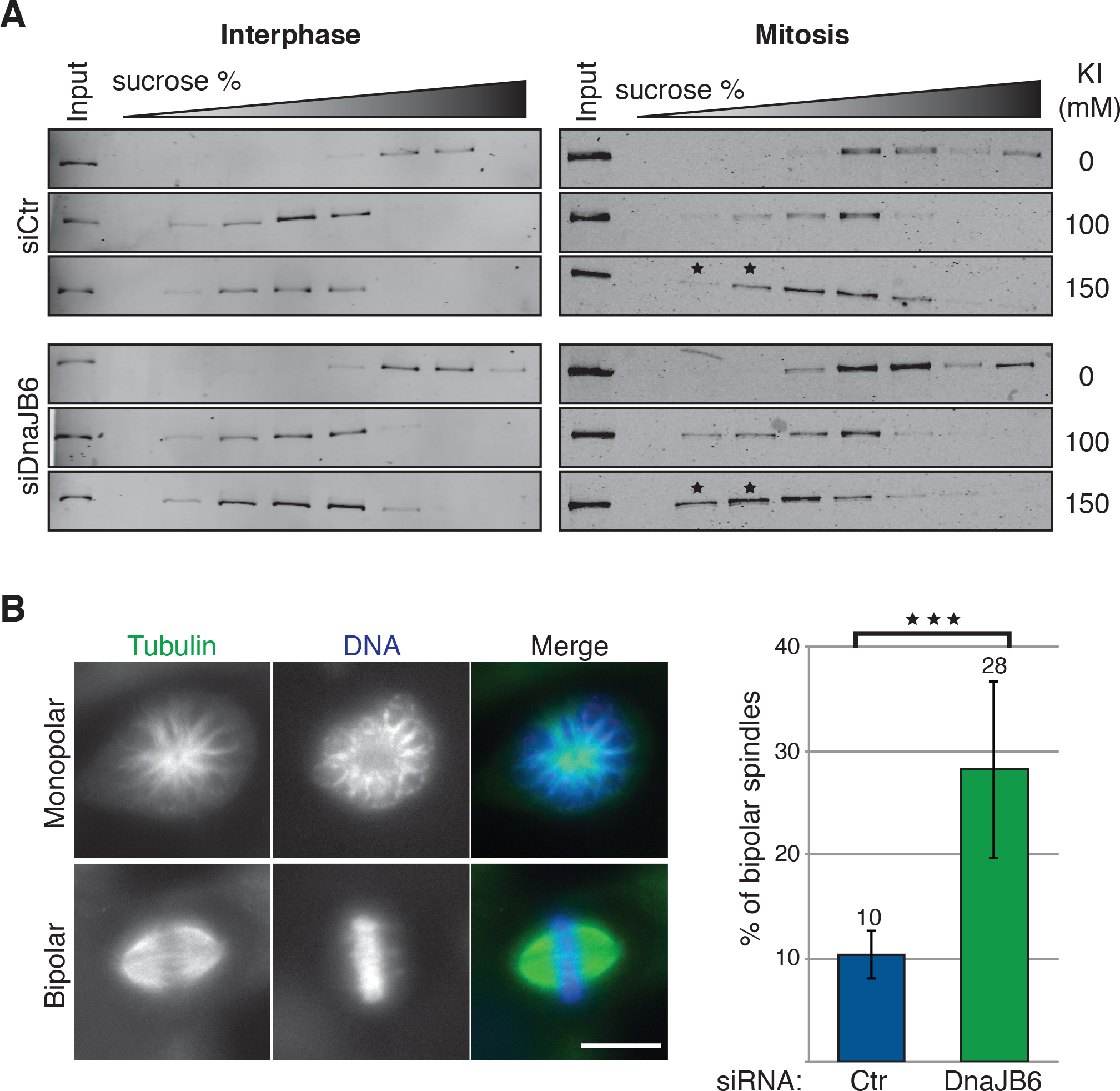
DnaJB6 is required for dynactin stability and dynein dependent force generation within the spindle. A. Western blot analysis showing the position of p150Glued in 8-20% sucrose density gradients from lysates of control and DnaJB6 silenced cells. Lysates were prepares form interphase (left) or mitotic (right) cells. KI was added to the lysates before running the gradients at the concentrations indicated on the left. A shift on the detection of p150 to lower fractions of sucrose is observed in mitotic DnaJB6 cell lysates treated with 150mM KI (right side lower band) compared to the mitotic control cells lysates with the same treatment (right side third band), highlighted using an asterisk.
B. DnaJB6 silencing rescues spindle bipolarity in STLC treated HeLa cells. Left: Immunofluorescence images of a representative monopolar and bipolar spindles in DnaJB6 silenced HeLa cells incubated with STLC. Tubulin is shown in green and DNA in blue. Scale bar, 10μm. Right: Bars graph showing the percentage of bipolar spindles in control or DnaJB6 silenced HeLa cells incubated with STLC. Data from three independent experiments in which 937 control and 1057 DnaJB6 silenced cells were analyzed. Three asterisks correspond to P<0.001(ANOVA test).

### DnaJB6 favors the generation of inward forces within the spindle

The dynactin complex is the major adaptor for dynein promoting its processivity and force generation (Cianfrocco et al., 2015; Grotjahn et al., 2018; Jha and Surrey, 2015). The phenotypes associated with the absence of DnaJB6 during mitosis are consistent with those described for dynein interference in mitosis, including spindle pole focusing defects and multipolarity (McKinley and Cheeseman, 2017; Raaijmakers and Medema, 2014). Moreover, our sucrose gradient results are compatible with DnaJB6 being required for the stability of the dynactin or dynein-dynactin complexes. This suggested that DnaJB6 silencing could impair dynein activity during mitosis. It has been previously shown that interfering with dynein activity rescues the formation of bipolar spindles in Eg5 deficient cells (Ferenz et al., 2009; Mitchison et al., 2005; Raaijmakers et al., 2013; Tanenbaum et al., 2008). We therefore asked whether DnaJB6 silencing could affect dynein force generation and rescue spindle bipolarity in STLC (S-trityl-l-cysteine, Eg5 inhibitor (Mayer et al., 1999)) treated HeLa cells. Control and DnaJB6 silenced HeLa cells were incubated with STLC for 2 hours, fixed and processed for immunofluorescence. As expected, control cells had a majority of monopolar spindles (88%). Interestingly, DnaJB6 silenced cells had a significant increase of bipolar spindles to 28% instead of 10% in the control cells (Fig 6B). To test whether DnaJB6 could function through Hsp70 for inward force regulation we tested whether a general HSP70 inhibitor (VER155008 (Massey et al., 2010; Schlecht et al., 2013)) could produce a similar bipolar spindle assembly rescue in STLC treated cells. We did not observe any rescue by inhibiting HSP70 indicating a direct role for DnaJB6 in the regulation of inward force production during spindle assembly (Fig S4A, B). Consistently we found a small but significant increase of the spindle length in DnaJB6 silenced HeLa cells (10.4μm) compared to control cells (9.9μm) (Fig S4C).

Altogether, our results suggest that DnaJB6 is required for dynein-dependent spindle inward forces generation during mitosis.

## Discussion

Here we showed that the nuclear protein DnaJB6 co-pulls down dynactin p150Glued in a RanGTP dependent way during mitosis. Moreover, the phenotypes observed upon changing the levels of DnaJB6 in cells and egg extracts are consistent with defects in dynein function. Our work therefore suggests a novel mechanism by which the small GTPase Ran in its GTP bound form regulates the activity of the dynein-dynactin complex during M-phase to promote inward force production and spindle pole focusing.

Two isoforms of DnaJB6 are present in human cells but only the long isoform localizes to the nucleus in interphase and interacts in a RanGTP dependent manner with P150Glued in M-phase. Silencing experiments reduced the levels of both the short and the long isoforms but the destabilization of the dynactin complex only occurred in mitosis suggesting that the interphase cytoplasmic short isoform plays no role in dynactin stabilization and function. Moreover, our data show that the long isoform of DnaJB6 fully rescues the phenotypes of the DnaJB6 depletion in egg extracts. Altogether, these data strongly suggest that the function in spindle assembly that we describe here is entirely specific for the long isoform of DnaJB6.

DnaJB6 is a member of the HSP40 protein family. The best characterized role of several members of this family is to act as co-chaperons of HSP70 proteins. Some members of the HSP70 family were previously shown to play a role in spindle assembly (Fang et al., 2016; O’Regan et al., 2015). Here we found that HSP70 inhibition does not rescue spindle bipolarity in Eg5 inhibited cells nor does it generate spindle pole focusing defects (data not shown). Moreover, the rescue experiments in egg extracts support a direct role for DnaJB6 in mitosis since the recombinant protein that rescued the depletion phenotype was added to the depleted extract only as the extract was sent into mitosis (Fig 4 and 5). Altogether, our data strongly suggest that the role of DnaJB6 in spindle assembly that we describe here is independent of HSP70. In agreement with this, HSP70 independent functions have been reported for some HSP40 family proteins including DnaJB6 (Hageman et al., 2010).

All the phenotypes observed in DnaJB6 silenced cells are compatible with an impairment of dynein function. Indeed, both spindle multipolarity and pole focusing defects may result from an unbalanced of forces within the spindle. The weakening of inward forces most probably result in dominant outward forces that may promote the disruption of the spindle pole integrity that may be associated with the formation of multipolar spindles (Maiato and Logarinho, 2014).

Dynein is a major minus-end directed motor that performs a large number of functions in the cell. During mitosis it is essential for centrosome separation, chromosome alignment, spindle pole focusing, spindle positioning and mitotic checkpoint silencing in addition to generating the inward forces that balance Eg5 (Kardon and Vale, 2009; Raaijmakers and Medema, 2014). The wide range of dynein functions raises the question of how this motor is regulated. Several interacting proteins were found to provide specificity for dynein localization and/or regulation (Cianfrocco et al., 2015). One of the best-characterized general regulators of dynein is dynactin, a large protein complex that participates in most dynein functions. Dynactin was shown to act as a dynein activator increasing its processivity and force production (Culver-Hanlon et al., 2006; Grotjahn et al., 2018; Kardon et al., 2009; King and Schroer, 2000; Ross et al., 2006; Tripathy et al., 2014).

We have shown that the long isoform of DnaJB6 pulls down the dynactin subunit p150Glued in a RanGTP-dependent manner during mitosis. This interaction increases resistance of the dynactin complex from KI induced disassembly specifically in mitosis. Surprisingly, NuMa and p150Glued are robustly recruited to the spindle poles in the absence of DnaJB6. Since our data from Eg5-inhibited cells strongly suggests that dynein force generation is defective in the absence of DnaJB6, the accumulation of NuMa and p150Glued to the poles may be a compensatory mechanism to promote their focusing.

In summary, we describe here the identification and functional characterization of DnaJB6 as a novel RanGTP-regulated protein required for microtubule organization during mitosis. Our data show that DnaJB6 interacts with the dynactin component p150Glued in a RanGTP-dependent manner during mitosis and regulates force generation by dynein-dynactin. Altogether our data suggest the existence of a mechanism by which RanGTP regulates dynein function and spindle organization.

## Materials and methods

### Hela cells culture, DnaJB6 silencing and drug treatments

Cells were grown at 37°C in a 5% CO2 humid atmosphere. HeLa cells were grown in DMEM 4,5g/L Glucose, with Ultraglutamine 1 (bioWhittaker, BE12-604F/U1), 10% FBS (102070-106, invitrogen) and 100units/ml penicillin and 100μg/ml streptomycin (15140-122, Invitrogen). Stable HeLa cells expressing H2B–eGFP/α- tubulin–mRFP were a gift from P. Meraldi (ETH, Zurich)(McAinsh et al., 2006) and were grown in the presence of 400μg/ml G418 and 20μg/ml puromycin. Cells were periodically tested for mycoplasma contamination.

Short RNA-mediated interference oligonucleotides targeting DnaJB6 (5’-CUAUGAAGUUCUAGGCGUG-3’) or scrambled (5’-CGUACGCGGAAUACUUCGAUU-3’) were transfected with lipofectamine RNAmax (13778-150, invitrogen) as detailed in the manufacturer protocol. 4μg of siRNA were used for each well of a six-well plate. All experiments were performed 48 hours after transfection.

HeLa cells were seeded on sterilized coverslips in 6 well plates, transfected with the selected siRNAs and incubated at 37°C for 48 hours.

For microtubule regrowth experiments, cells were incubated for 3 hours at 37°C with 2μM nocodazole (Sigma, M1404). Nocodazole was then washed out by 3 washes with 2ml of pre-warmed PBS1X and one of media at 370C. Cells were incubated for different periods in a 370C incubator, fixed in −20°C methanol for 10 minutes and processed for IF using the DM1A antibody.

To perform K-Fibers stability tests, cells were incubated in cold K-fiber medium (L15 medium (Sigma) supplemented with 20mM HEPES (Sigma) at pH 7.3) on ice for 10, 20 or 30 minutes before fixation in −20°C methanol for 10 minutes and processed for IF with the DM1A antibody to visualize the microtubules.

To measure K-Fiber length, cells were incubated with 10μM STLC (Sigma, 164739) during 3 hours at 37°C and placed on ice with cold K-fiber medium containing 10μM STLC for 7 minutes before fixation in −20°C methanol for 10 minutes and for IF with the DM1A antibody to visualize the microtubules. Images were analyzed with FIJI. The length of individual K-fibers was measured in three independent experiments.

For the spindle bipolarization rescue experiment, control and DnajB6 silenced cells were incubated for 2 hours at 37°C with 2μM STLC (Sigma, 164739), fixed in −20°C methanol and processed for IF using the DM1A or anti-tubulin-β antibodies.

### Cloning of Xenopus DnaJB6

Xenopus mRNA was obtained from CSF *Xenopus leavis* egg extracts using TRIzol reagent (15596-026, thermofisher) according to manufactures instructions. cDNA was generated by RT-PCR using an oligo-dT coupled with an adaptor sequence (5’-GGCCACGCGTCGACTAGTAC +17 “T”-3’). The 3’ end sequence of the mRNA was obtained by PCR using the Phusion High-Fidelity DNA Polymerase (F530S; thermofisher) according to the manufacturer instructions, using a primer with the adaptor sequence and another corresponding to the central part of the xDnaJB6-S (NM_001095775.1) annotated sequence that is conserved between the long and the short isoforms in human (5’-GGAGGTTTCCCTGCCTTTGGCCC-3’). In a second step, another PCR was performed on the Xenopus egg cDNA using a reverse primer corresponding to the 3’end of the long isoform previously obtained and a forward primer containing the annotated start codon from xDnaJB6-S (fwd 5’-ATGGTGGAGTATTACGAAGTTTTGGGAGTCC-3’; rev 5’ TTAGTAGATTGGTTTGGAAGACTTCTTTTTCTTG-3’). The sequence for xDnaJB6 long isoform will be deposited into GenBank database.

### Protein production

The full length xDnaJB6-L was cloned in pGex-4T-2 and pMal-c2 vectors. MBP-xDnaJB6 was expressed in the E.coli BL21/RIL strain by induction with 0,5 mM IPTG for 4 hours at 25°C. The protein was purified on amylose resin beads (E8021S, NEB) according to manufacturer instructions. The purified protein was stored in 20mM Tris-HCl pH 7.4, 200mM NaCl at −80°C.

GST-xDnaJB6 was expressed in the E.coli BL21pLys strain by incubation with 1mM IPTG ON at 20°C. For protein purification glutathione sepharose beads (17-5132-01, GE healthcare) were used according to manufactures instructions. Protein was kept in 50mM tris-HCl PH 8.0, 200mM KCL at −80°C.

### Antibodies

Affinity purified antibodies against hDnaJB6 and GST were previously described (Rosas-Salvans et al., 2018). Antibodies against xDnaJB6 were obtained by rabbit immunization with MBP-xDnaJB6-L and affinity purified. The working concentrations were the following: anti-hDnaJB6 (IF-5μg/ml; WB-1μg/ml), anti-xDnaJB6 (WB-1μg/ml).

The following primary commercial antibodies were used: DM1A (Sigma, T6199; IF and WB-1μg/ml), anti-tubulin-β (Abcam, ab6046; IF-1:200), anti-p150Glued (BD transduction laboratories, 610474; IF-5μg/ml; WB-0,5μg/ml), anti-centrin (Millipore, 04-1624; IF-1:1000), anti-NUMA (Calbiochem, NA09L; IF-1:500), anti-Importin-β/NTF97 (Abcam; ab36775-50; WB-1:500) and generic IgGs (Sigma, I5006). The following secondary antibodies were used: Alexa Fluor 488 Goat, anti-mouse (A11017; Invitrogen), Alexa Fluor 488 goat, anti rabbit (A11034; Invitrogen), Alexa Fluor 568 anti-mouse (A11031; Invitrogen) and Alexa Fluor 568 goat, anti-rabbit (A11036; life technologies S.A) all at 0,2μg/ml final concentration. For Western blots, the following antibodies were used: anti-mouse 800 (926-32212; Licor), goat anti-rabbit 800 (10733944; Fisher Scientific), goat anti-Rabbit Alexa Fluor 680 (A-21109; Invitrogen) and goat anti-mouse Alexa Fluor 680 (A21058; Life technologies) diluted at 1:10000.

### Xenopus egg extracts

All the experiments involving animals were performed according to standard protocols approved by the ethical committee of the Parc de Recerca Biomèdica de Barcelona. *Xenopus laevis* frogs (males and females) were purchased from Nasco and used at an age between 1 and 3 years. CSF arrested egg extracts were prepared and used to performed cycled spindle assembly assays as previously described (Desai et al., 1999).

For depletion experiments, protein A-conjugated Dynabeads 280 (10002D, invitrogen) were coated with α-xDnaJB6 or generic IgGs (9μg of antibody/30μl of beads) and incubated in incubated in freshly prepared egg extract for 30min on ice. Beads were recovered and protein depletion efficiency tested by western blot analysis. For add back experiments, 0,017μM (estimated endogenous concentration of DnaJB6 in the egg extract) of MBP-xDnaJB6-L or MBP were added to the depleted egg extracts. To look at the effects of increasing the concentration of DnaJB6 we added 1μM of MBP or MBP-xDnaJB6-L to egg extracts.

*Xenopus laevis* frogs were purchased from Nasco and used at the age of 1 to 3 years. Animals were housed at the animal facility of the PRBB and handled following protocols approved by the ethical committee of the Parc de Recerca Biomèdica de Barcelona (PRBB).

### Pull-down experiments from egg extracts

Protein A-conjugated Dynabeads 280 (10002D, invitrogen) were coated with α-GST or α-MBP (9μg of antibody/30μl of beads), and incubated with recombinant proteins (GST; GST-xDNAJB6-L; MBP or MBP-xDnaJB6-L) Protein coated beads were then incubated in CSF egg extract (30μl of beads/75μl of EE) for 15 minutes at 20°C followed by 30 minutes on ice. Recombinant RanQ69L-GTP was added at 15μM. Beads were recovered and washed three times in CSF-XB (for checking importin interaction) or three times in CSF-XB and one in PBS-NP40 (for checking p150Glued interaction). Proteins were eluted from the beads by incubation in sample buffer for 10 minutes RT and run on SDS-PAGE before Western blotting.

### Immunofluorescence

Cells grown on coverslips were fixed in −20°C methanol for 10 minutes. They were blocked and permeabilized for 30 minutes in IF-buffer (PBS1X, 0.1% tritonX100, 0.5% BSA). Primary antibodies diluted in IF-buffer were then applied for 1hour at room temperature. Samples were washed 3 times in IF buffer and secondary antibodies and Hoechst (Hoechst 33342; ref: H3570; Invitrogen; added at 1:1000) were applied for 45 minutes. The coverslip were washed with IF buffer and twice with PBS before mounting in Mowiol.

For anti-DnaJB6 immunostaining cells were pre-extracted for 6 seconds in BRB80-1X, 0,5% triton and 1mM DSP. Cells were then fixed in PFA 4% for 7 minutes and then processed for IF as described.

Spindles assembled in egg extracts were centrifuged onto coverslips as previously described, fixed in −20°C methanol for 10 minutes and processed for IF following the above protocol.

### Sucrose gradients

Sucrose gradients (8 to 20%) experiments were performed as described in (Jones et al. 2014). For preparing interphase cell lysates cells were collected in 5ml of PBS using a scraper. For preparing mitotic lysates, cells were incubated for 15 hours in 2μM nocodazole, washed 3x with PBS and once with medium (20ml). They were then incubated at 37°C for 45 minutes when the majority of the cells were in metaphase. Cells were recovered by shake off. To prepare the lysates, cells were washed twice in BRB80-1X, centrifuged at 600g for 5 minutes. The pellets were resuspended in 500μl of lysis buffer (BRB80-1X, 0.5% Triton-X100 with protease inhibitors) for 15 minutes on ice and subsequently centrifuged at 13200rpm (in a 5415 R Eppendorf centrifuge) for 10 min at 4°C. The protein concentration of the supernatants was measured by Bradford. 100μl of control or DnaJB6 silenced cell lysates were incubated with potassium iodide (0, 100 or 150 μM) and incubated for 1 hour on ice. 45μl of each sample was added to the top of sucrose gradients in centrifuge tubes previously prepared at room temperature. Tubes were centrifuged at 38000rpm for 5 h at 4°C in a SW-55Ti rotor with an ultracentrifuge Beckman optima L-100K. 75μl fractions were recovered from the top and diluted in sample buffer. Samples were then loaded on SDS PAGE gels followed by a Western blotting.

Sucrose gradients were prepared in 800 μL (0.8 mL) Ultra-Clear Centrifuge Tubes, 5 × 41 mm (Part Number 344090) and specific adaptors used for centrifugation (Adapter, Split, Delrin, Tube, 5mm diameter (qty. 2 halves, product number 356860). Sucrose solutions were prepared in BRB80 containing protease inhibitors (1/4000), 0,1mM ATP, 1mM DTT, sucrose and KI at the selected concentrations (0, 100 or 150μM KI). The gradient was prepared by overlaying manually the different sucrose solutions (2% difference of sucrose between layers, 7 layers from 20 to 8% of sucrose) freezing between each layer addition. The gradient were kept at −20°C and warmed up at RT for one hour before use.

### Microscopy

Non-synchronized HeLa cells stably expressing H2B–eGFP/α-tubulin–mRFP were imaged every 4min for 18 hours with a 40X objective on a Zeiss Cell Observer HS inverted microscope equipped with a 40X objective, a Zeiss AxioCam MrX camera and a temperature, humidity and CO2 control systems. Images were processed and analyzed with FIJI.

Immunofluorescence images were obtained with an inverted DMI-6000-B Leica wide-field fluorescent microscope equipped with 40X and 63X objectives and a Leica DFC 360FX camera. Confocal images were obtained on a Leica TCS SPE microscope equipped with a 63X objective and the following laser lines: 405 (DAPI, Hoechst), 488 (GFP, FITC), 532 (mRFP, DsRED, Cy3, TRITC, TexasRed), 635 (Cy5, DRAQ5).

### Fluorescence intensity profile quantifications

Images taken with a 63X objective on an inverted DMI-6000-B Leica wide-field fluorescent microscope, equipped with a Leica DFC 360FX camera, were analyzed using the FIJI program as follows. Spindles were individually rotated in order to orient them with an horizontal pole-to-pole axis. A rectangle of a fixed size containing both spindle poles and centered on the metaphase plate was drawn on each image. The average pixel intensity was measured for sequential vertical pixel lines, obtaining a list of intensities, which was exported to an independent file. The intensity lists for all the cells analyzed in any given condition were analyzed using the software Prism and plotted together. A line graph was generated with a line showing the average intensities in each vertical pixel line and the standard deviations. In order to compare the protein intensities at the spindle poles, the length of the spindles was normalized by subtracting the central spindle values of the larger spindles, when necessary. Protein intensities were normalized using the intensity of the tubulin (obtained with the same method). The graphs obtained for DnaJB6 silenced cells and control cells were compared with the ANOVA test to obtain the significance of the differences between the two conditions.

### Statistical analysis

PRISM software was used for the statistical analysis. Depending on the samples the ANOVA test, the Mann-whitney test or the Fisher’s exact test was used. Normality distribution was tested (D’Agostino-pearson omnibus normality test) for each sample when applied and equality of variances between samples compared conditions was tested when necessary. Sample size was determined based on previous work done in the laboratory. For quantifications, the investigator was blinded to the sample allocation at the moment of counting.

### Data availability

Data from this work are available upon request.

## Acknowledgements

We thank S. Meunier for discussions and initial supervision. We thank N. Mallol and L. Avila for excellent technical support. We thank the CRG microscopy facility for technical support. We acknowledge the Spanish Ministry of Economy, Industry and Competitiveness (MEIC) to the EMBL partnership and support of the Spanish Ministry of Economy and Competitiveness, ‘Centro de Excelencia Severo Ochoa’ as well as support of the CERCA Programme / Generalitat de Catalunya. M.R. was supported by the FPI fellowship from the Spanish Ministry of Economy BES-2013-064601. Work in the Vernos lab was supported by grants from the Spanish Ministry of Economy (MINECO) I+D grants BFU2012-37163 and BFU2015-68726-P.

## Author contributions

MR performed the experiments, analyzed the data, prepared figures and wrote the manuscript.

IV designed the project, prepared figures and wrote the manuscript.

## Materials and correspondence

Material and correspondence should be addressed to I.V.

