## Supplementary material for "DnaJB6 is a RanGTP-regulated protein involved in dynein-dependent microtubule organization during mitosis": suplementary figures

**Supplementary figure legends**

**FIGURE S1**

**A)** Western blot analysis of control and DnaJB6 silenced cells. Cells were lysed 48 hours post-transfection and 30 µg of protein were loaded per lane. DnajB6 silencing efficiency was very high: 90% for DnaJB6-L and more than 90% for DnaJB6-S.

**B)** DnaJB6 silenced cells have ectopic microtubule clusters in metaphase.

Left: Immunofluorescence images of a representative DnaJB6 silenced metaphase cell with an ectopic microtubule cluster (white arrow). In the merge (upper image) tubulin is shown in green and DNA in blue. Scale bar, 10µm.

Right: Graph showing the percentage of bipolar spindles containing ectopic microtubule clusters in non synchronized control and DnaJB6 silenced HeLa cells. 182 control and 192 DnaJB6 silenced cells in metaphase were examined. Three asterisks correspond to $P<0.001$ (Fisher exact test).

**C)** Box and whiskers plot of the number of microtubule asters formed in mitotic control and DnaJB6 silenced cells in a MT regrowth assay. Cells were fixed at the indicated time-points after nocodazole washout. Data from three independent experiments (total sample size: control cells: 350, 328, 310, 333, 317 control and DnaJB6 silenced cells: 353, 301, 330, 354 and 312). Boxes show values between the 25th and the 75th percentiles, the black lines correspond to the median and + to the mean. Whiskers extend from the 10th to the 90th percentile and dots correspond to outliers. Three asterisks correspond to P-value<0.0001 (Mann-Whitney test).

**FIGURE S2**

**A)** Western blot analysis of control and xDnaJB6 depleted Xenopus egg extracts without (-) or with (+) recombinant MBP-xDnaJB6-L. The depletion was very efficient and the recombinant protein was added at close to endogenous concentrations. 1µl of egg extract was loaded per lane. The endogenous and recombinant proteins were detected with an Anti-xDnaJB6 antibody.

**FIGURE S3**

**A)** Quantification of the p150Glued and tubulin fluorescence intensities measured along the spindle in control (blue) and DnaJB6 silenced HeLa cells (green). Protein fluorescence intensities were analyzed using FIJI. The relative positions along the spindle are indicated with arbitrary units. Mean values and the standard error of the mean are shown. P150Glued fluorescence intensity is significantly increased at the spindle poles in DnaJB6 silenced cells. The significance was calculated after normalization of the p150Glued signal on the tubulin signal ($P<0.05$, two tailed ANOVA test). The measurements were obtained from 47 (control) and 37 (DnaJB6 silenced) metaphase spindles.

**B)** Quantification of the NuMA and tubulin fluorescence intensities measured along the spindle in control (blue) and DnaJB6 silenced HeLa cells (green). Protein fluorescence intensities were analyzed using FIJI. The relative positions along the spindle are indicated with arbitrary units. Mean values and the standard error of the mean are shown. NuMA fluorescence intensity is significantly increased at the spindle poles in DnaJB6 silenced cells. The significance was calculated after normalization of the p150Glued signal on the tubulin signal ($P<0.05$, two tailed ANOVA test). The measurements were obtained from 56 (control) and 36 (DnaJB6 silenced) metaphase spindles.

**FIGURE S4**

**A)** Graph showing the percentage of bipolar spindles in control and HSP70 inhibited HeLa cells treated with STLC. Cells were incubated with the HSP70 inhibitor Ver155008 at 5µM final concentration. A significant reduction of the percentage of bipolar spindles was detected in HSP70 inhibited cells. Data from two independent experiments monitoring 544 control and 549 HSP70 inhibited cells. Two asterisks correspond to p-value<0.01 (two tailed ANOVA test).

**B)** Graph showing the percentage of bipolar spindles in control and HSP70 inhibited HeLa cells treated with STLC. Cells were incubated with the HSP70 inhibitor Ver155008 at 40µM. Data from two independent experiments monitoring 450 control and 472 HSP70 inhibited cells. No significant differences were detected between the two conditions (two tailed ANOVA test).

**C)** Box and whiskers plot showing the spindle lengths obtained in control and DnaJB6 silenced HeLa cells. Boxes show values between the 25th and the 75th percentiles, the black line corresponds to the median and + to the mean. Whiskers extend from the 10th to the 90th percentile and dots correspond to outliers. The spindle length of 267 control and 294 DnaJB6 silenced cells from three independent experiments was measured. A statistically significant increase of the length was detected in DnaJB6 silenced cells Three asterisks correspond to p-value<0.0001 (Mann-Whitney test).

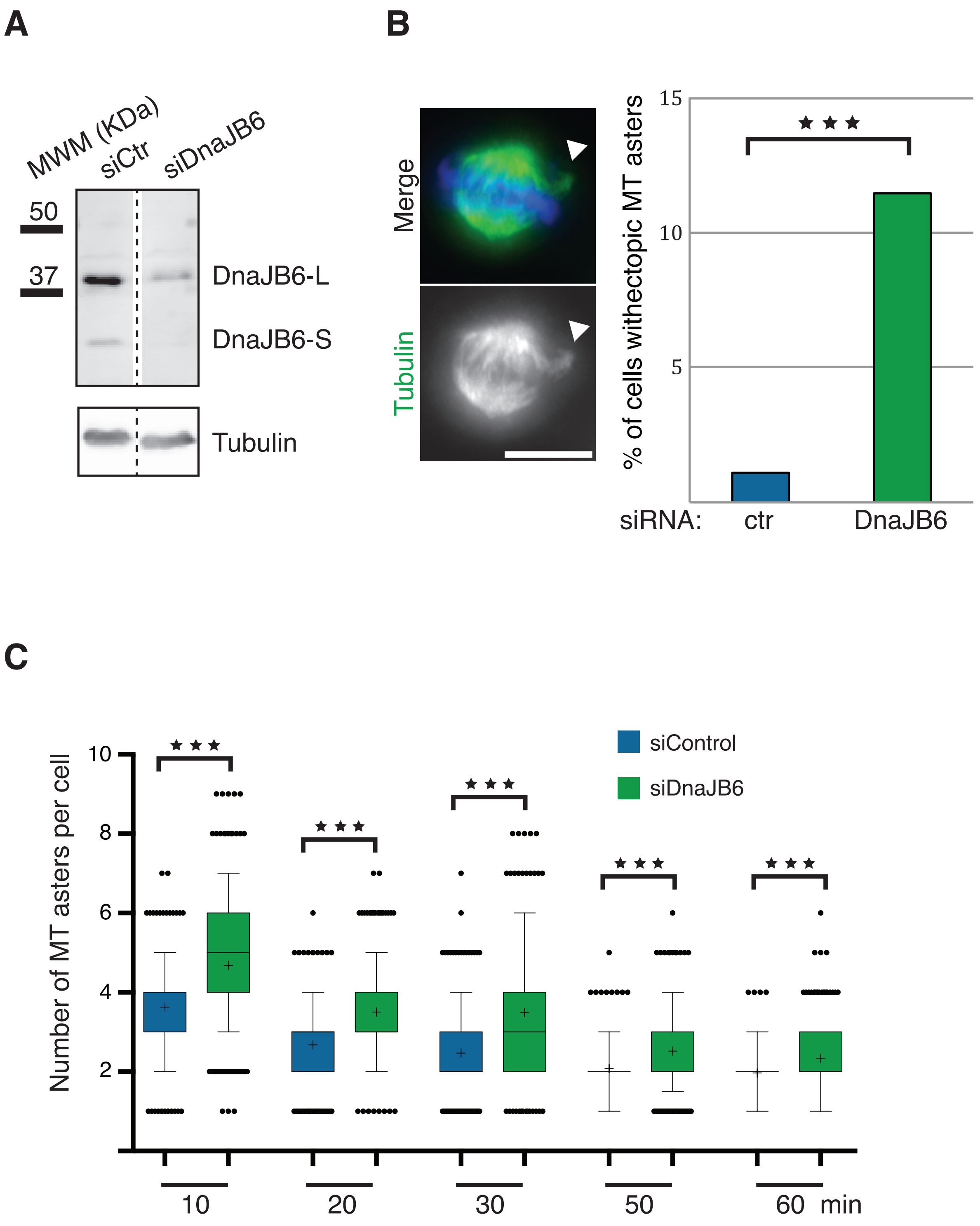
A
B
C
MWM (KDa)
siCtr
siDnaJB6
50
37
DnaJB6-L
DnaJB6-S
Tubulin
Merge
Tubulin
% of cells withectopic MT asters
15
10
5
siRNA:
ctr
DnaJB6
Number of MT asters per cell
siControl
siDnaJB6
10
8
6
4
2
10
20
30
50
60 min

Figure S1

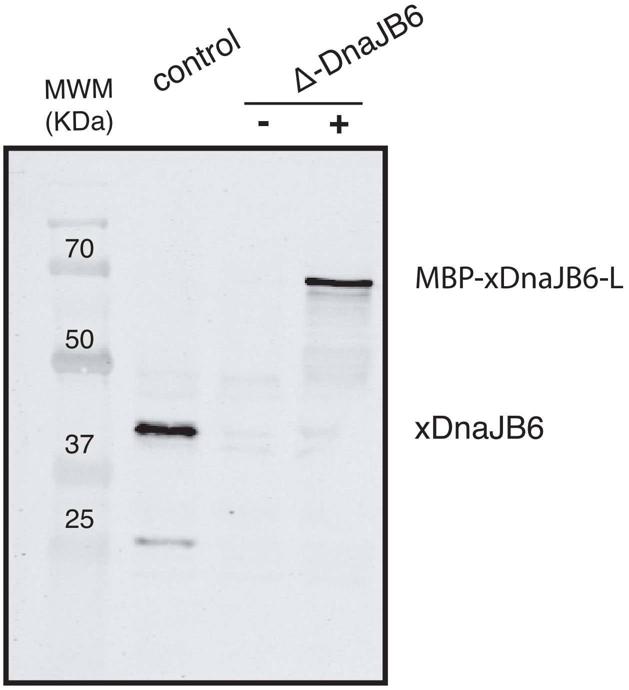


**Figure S2**

## A

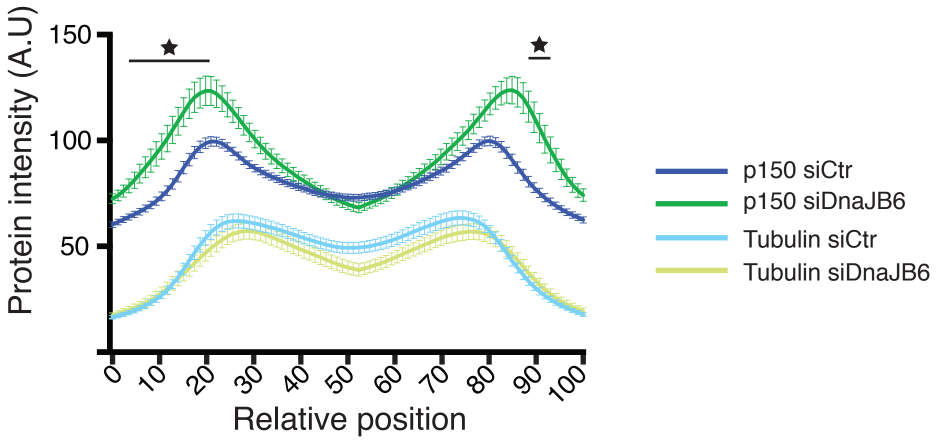

150
100
50
Protein intensity (A.U)
0
10
20
30
40
50
60
70
80
90
100
Relative position
p150 siCtr
p150 siDnaJB6
Tubulin siCtr
Tubulin siDnaJB6


## B

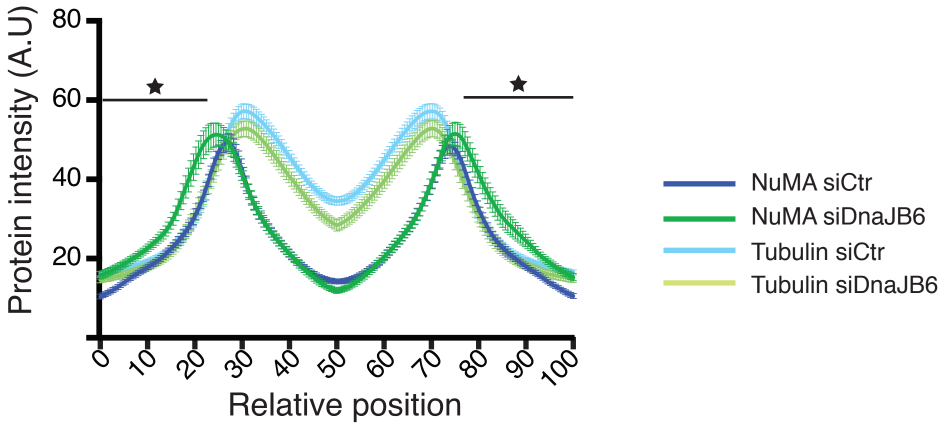

80
60
40
20
Protein intensity (A.U)
0
10
20
30
40
50
60
70
80
90
100
Relative position
NuMA siCtr
NuMA siDnaJB6
Tubulin siCtr
Tubulin siDnaJB6


### Figure S3

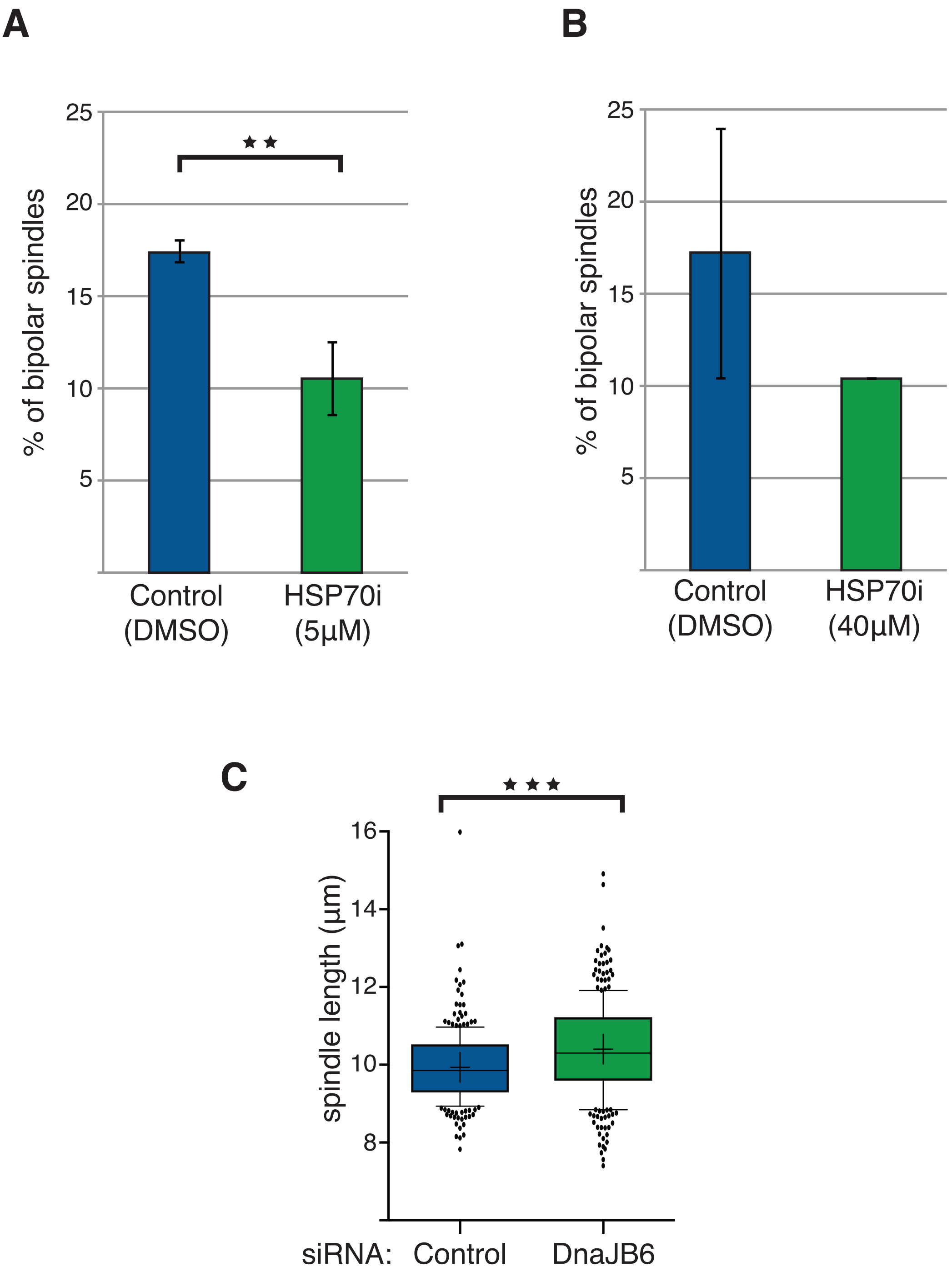

A
25
20
15
10
5
% of bipolar spindles
Control
(DMSO)
HSP70i
(5µM)
B
25
20
15
10
5
% of bipolar spindles
Control
(DMSO)
HSP70i
(40µM)
C
16
14
12
10
8
spindle length (µm)
siRNA:
Control
DnaJB6


**Figure S4**
